# Cell specific eQTL analysis without sorting cells

**DOI:** 10.1101/002600

**Authors:** Harm-Jan Westra, Danny Arends, Tõnu Esko, Marjolein J. Peters, Claudia Schurmann, Katharina Schramm, Johannes Kettunen, Hanieh Yaghootkar, Benjamin P. Fairfax, Anand Kumar Andiappan, Yang Li, Jingyuan Fu, Juha Karjalainen, Mathieu Platteel, Marijn Visschedijk, Rinse Weersma, Silva Kasela, Lili Milani, Liina Tserel, Pärt Peterson, Eva Reinmaa, Albert Hofman, André G. Uitterlinden, Fernando Rivadeneira, Georg Homuth, Astrid Petersmann, Roberto Lorbeer, Holger Prokisch, Thomas Meitinger, Christian Herder, Michael Roden, Harald Grallert, Samuli Ripatti, Markus Perola, Andrew R. Wood, David Melzer, Luigi Ferrucci, Andrew B. Singleton, Dena G. Hernandez, Julian C. Knight, Rossella Melchiotti, Bernett Lee, Michael Poidinger, Francesca Zolezzi, Anis Larbi, De Yun Wang, Leonard H. van den Berg, Jan H. Veldink, Olaf Rotzschke, Seiko Makino, Timouthy M. Frayling, Veikko Salomaa, Konstantin Strauch, Uwe Völker, Joyce B. J. van Meurs, Andres Metspalu, Cisca Wijmenga, Ritsert C. Jansen, Lude Franke

**Author notes:** These authors contributed equally to this work (Harm-Jan Westra, Danny Arends). These authors jointly directed this work (Ritsert C. Jansen, Lude Franke).

## Abstract

Expression quantitative trait locus (eQTL) mapping on tissue, organ or whole organism data can detect associations that are generic across cell types. We describe a new method to focus upon specific cell types without first needing to sort cells. We applied the method to whole blood data from 5,683 samples and demonstrate that SNPs associated with Crohn's disease preferentially affect gene expression within neutrophils.

## Introduction

In the past seven years, genome-wide association studies (GWAS) have identified thousands of genetic variants that are associated with human disease (Hindorff et al. 2009). The realization that many of the disease-predisposing variants are non-coding and that single nucleotide polymorphisms (SNPs) often affect the expression of nearby genes (i.e. *cis*-expression quantitative trait loci; *cis*-eQTLs) (Fehrmann et al. 2011) suggests these variants have a predominantly regulatory function. Recent studies have shown that disease-predisposing variants in humans often exert their regulatory effect on gene expression in a cell-type dependent manner (Brown et al. 2013; Fairfax et al. 2012; Fu et al. 2012). However, most human eQTL studies have used sample data obtained from mixtures of cell types (e.g. whole blood) or a few specific cell types (e.g. lympoblastoid cell lines) due to the prohibitive costs and labor required to purify subsets of cells from large samples (by cell sorting or laser capture micro-dissection). In addition, the method of cell isolation can trigger uncontrolled processes in the cell, which can cause biases. In consequence, it has been difficult to identify in which cell types these disease-associated variants exert their effect. Here we describe a generic approach that deepens our interpretation of GWAS data to the level of individual cell types (Figure 1). Our strategy was to: (i) collect gene expression data from an entire tissue; (ii) predict the abundance of its constituent cell types (i.e. the cell counts) by using expression levels of genes that serve as proxies for these cell types; (iii) run an association analysis with a term for interaction between the SNP marker and the proxy for cell count to detect cell-type-mediated or -specific associations, and (iv) test whether known disease associations are enriched for SNPs that show the cell-type-mediated or -specific effects on gene expression (i.e. eQTLs).

**Figure 1.**
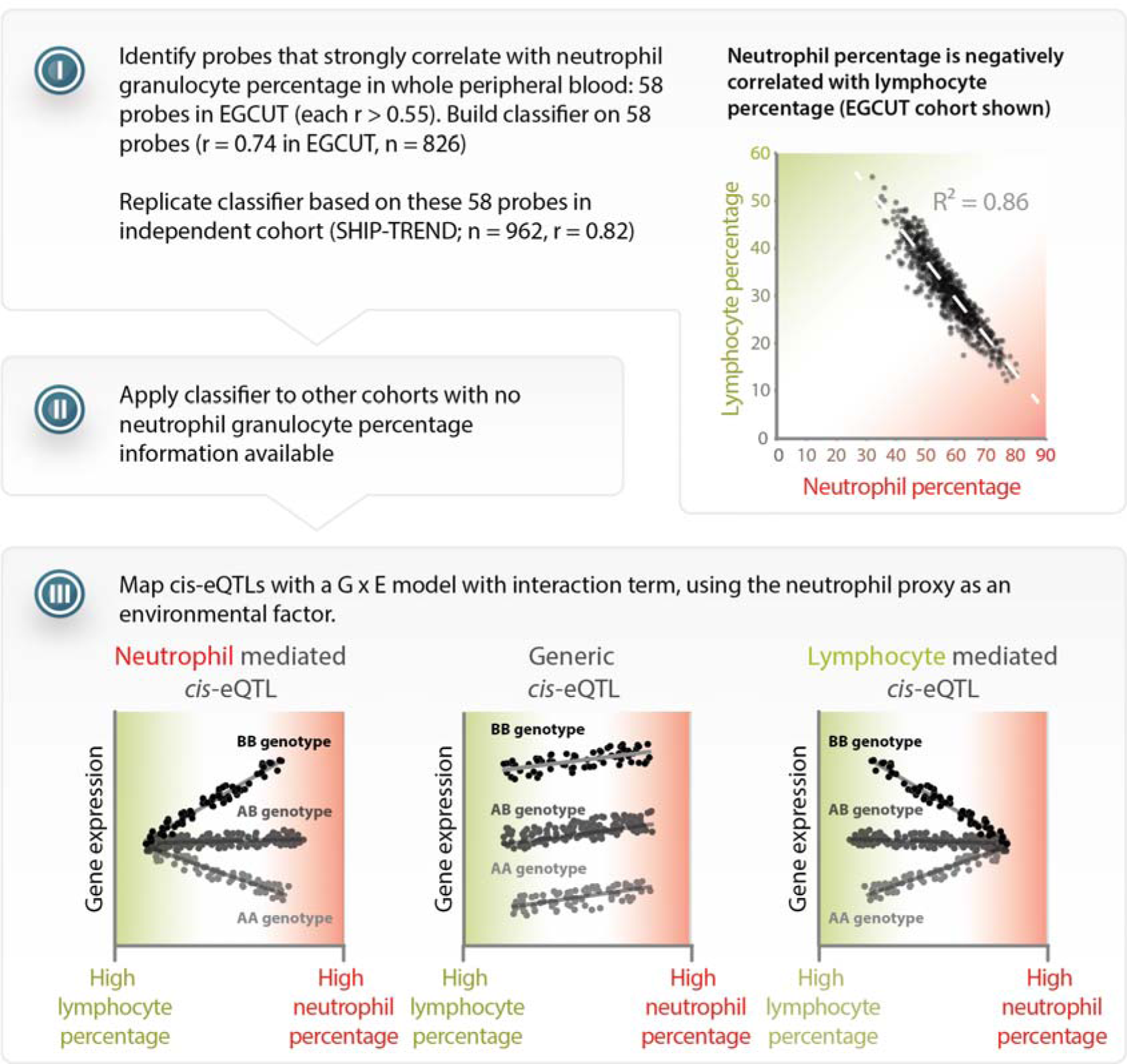
Method overview, power analysis and validation. A) Overview of the method to detect cell-type-mediated *cis*-eQTL in compound tissues. I) Starting with a dataset that has cell count measurements, determine a set of probes that have a strong positive correlation to the cell count measurements. Calculate the correlation between these specific probes in the other datasets, and apply principal component analysis to combine them into a single proxy for the cell count measurement. II) Apply the prediction to other datasets lacking cell count measurements. III) Use the proxy as a covariate in a linear model with an interaction term in order to distinguish cell-type-mediated from non-cell-type-mediated eQTL effects.

## Results

We applied this method to 5,863 unrelated, whole blood samples from seven cohorts: EGCUT(Metspalu 2004), InCHIANTI (Tanaka et al. 2009), Rotterdam Study (Hofman et al. 2013), Fehrmann (Fehrmann et al. 2011), SHIP-TREND (Völzke et al. 2011), KORA F4 (Mehta et al. 2013), and DILGOM (Inouye et al. 2010). Blood contains many different cell types that originate from either the myeloid (e.g. neutrophils and monocytes) or lymphoid lineage (e.g. B-cells and T-cells). Even though neutrophils comprise ∼62% of all white blood cells, no neutrophil eQTL data have been published to date, because this cell type is particularly difficult to purify or culture in the lab (Grisham et al. 1985).

For the purpose of illustrating our new cell-type specific analysis in the seven whole blood cohorts, we focused on neutrophils. Direct neutrophil cell counts and percentages were only available in the EGCUT and SHIP-TREND cohorts, requiring us to infer neutrophil percentages for the other five cohorts. We used the EGCUT cohort as a training dataset to identify a list of 58 Illumina HT12v3 probes that correlated positively with neutrophil percentage (Spearman's correlation coefficient R > 0.55). We then summarized the gene expression levels of these 58 individual probes into a single neutrophil percentage estimate, by applying principal component analysis (PCA) and using the first principal component (see Figure 2 for confirmation of the accuracy of prediction in the SHIP-TREND cohort; Spearman R = 0.81). We then used this procedure in the other cohorts to predict the neutrophil percentage.

**Figure 2.**
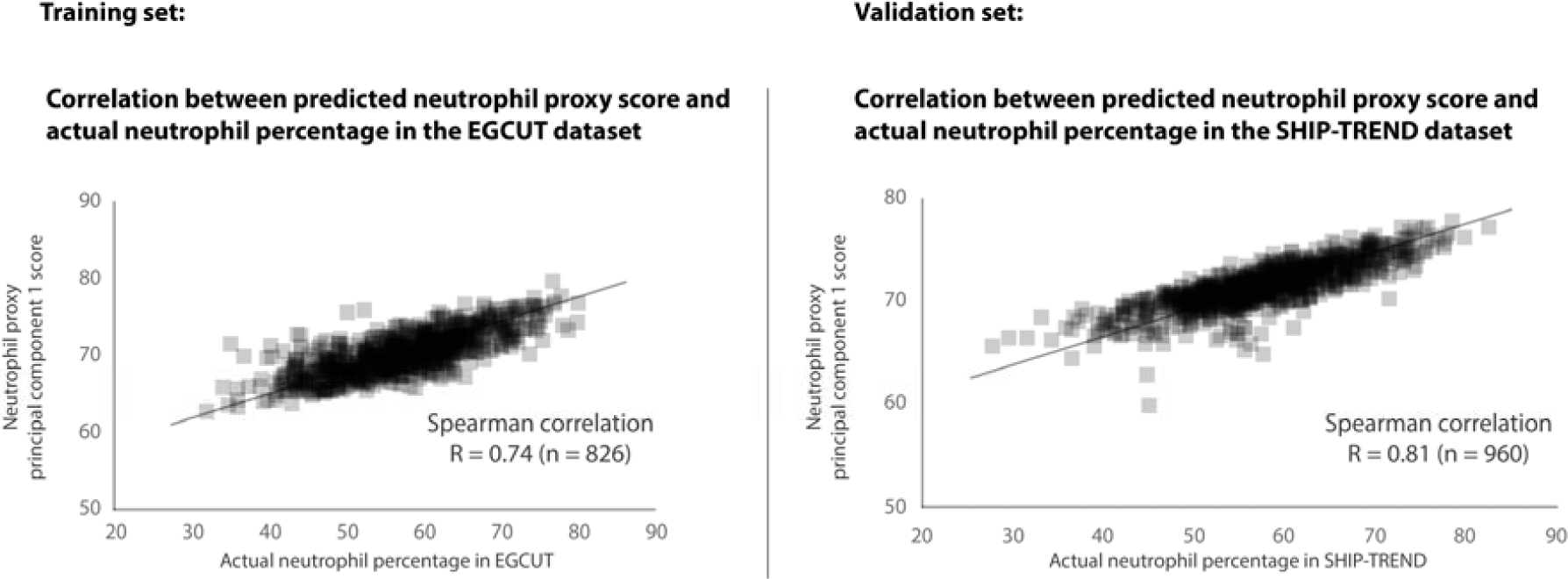
Validation of neutrophil proxy. There is a strong correlation between the neutrophil proxy and the actual neutrophil percentage measurements in the training dataset (EGCUT, r = 0.74). Validation of neutrophil prediction in the SHIP-TREND cohort shows a strong correlation (r = 0.81) between the neutrophil proxy and actual neutrophil percentage measurements in this dataset.

Here we limit our analysis to 13,124 previously discovered *cis*-eQTLs (Westra et al. 2013), (although a genome-wide application of our method might result in the detection of additional cell-type-specific *cis*-eQTLs, that we might have missed by assuming a generic effect across cell types). We performed the eQTL association analysis with a term for interaction between the SNP marker and the proxy for cell count within each cohort, followed by a meta-analysis (weighted for sample size) across all the cohorts. We identified 1,117 *cis*-eQTLs with a significant interaction effect (8.5% of all *cis*-eQTLs tested; false discovery rate (FDR) < 0.05; 1,037 unique SNPs and 836 unique probes; Supplementary Tables 1 and 2). Out of the total number of *cis*-eQTLs tested, 909 (6.9%) had a positive direction of effect, which indicates that these *cis*-eQTLs show stronger effect sizes in neutrophils (‘neutrophil-mediated *cis*-eQTLs’; 843 unique SNPs and 692 unique probes). Another 208 (1.6%) had a negative direction of effect (196 unique SNPs and 145 unique probes), indicating a stronger *cis*-eQTL effect size in lymphoid cells (‘lymphocyte-mediated *cis*-eQTLs’; since lymphocyte percentages are strongly negatively correlated with neutrophil percentages, Figure 1). Overall, the directions of the significant interaction effects were consistent across the different cohorts, indicating that our findings are robust (Supplementary Figure 1).

We validated the neutrophil- and lymphoid-mediated associations we detected in six small, purified cell-type gene expression datasets that had not been used in our meta-analysis. We generated new eQTL data from two lymphoid cell types (CD4+ and CD8+ T-cells) and one myeloid cell type (neutrophils, see online methods) and used previously generated eQTL data on two lymphoid cell types (lymphoblastoid cell lines and B-cells) and another myeloid cell type (monocytes, Supplementary Table 3). As expected, compared to *cis*-eQTLs without a significant interaction term (‘generic *cis*-eQTLs’, n = 12,007) the 909 neutrophil-mediated *cis*-eQTLs did indeed show very strong *cis*-eQTL effects in both of the myeloid datasets (Wilcoxon P-value ≤ 4.9 × 10^−31^), and small effect sizes in the lymphoid datasets. Conversely, the 208 lymphoid-mediated *cis*-eQTLs had a pronounced effect in each of the lymphoid datasets (Wilcoxon P-value ≤ 7.8 × 10^−14^; Figure 3A), while having small effect sizes in the myeloid datasets. These results indicate that our method is able to reliably predict whether a *cis*-eQTL is mediated by a specific cell type. Unfortunately, the cell type that mediates the *cis*-eQTL is not necessarily the one in which the *cis*-gene has the highest expression (Figure 3B), making it impossible to identify of cell-type-specific eQTLs directly on the basis of expression levels.

**Figure 3.**
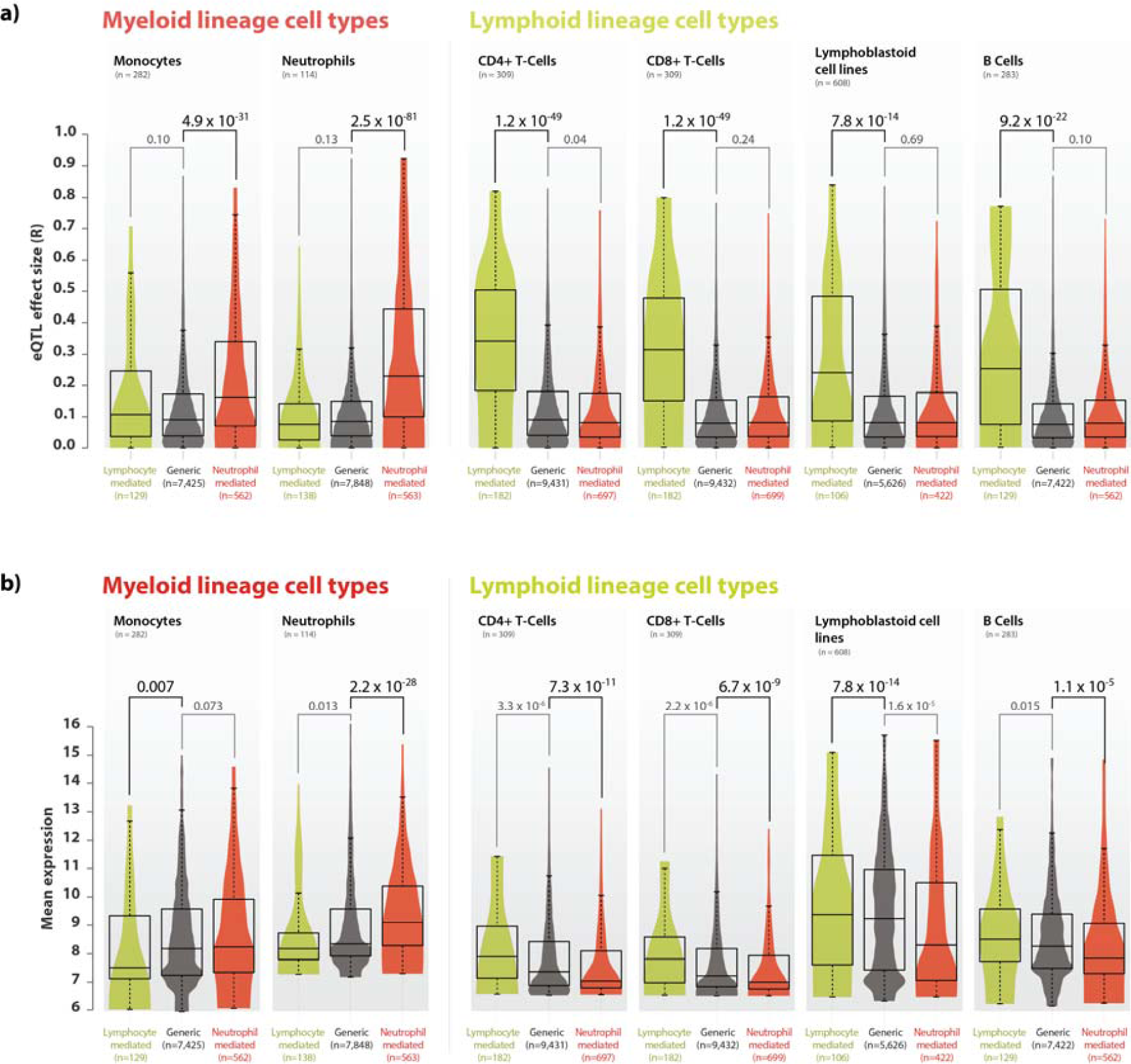
Validation of neutrophil and lymphoid specific <u>*cis*</u>-eQTLs in purified cell type eQTL datasets. A) We validated the neutrophil- and lymphoid-mediated *cis*-eQTL effects in four purified cell type datasets from the lymphoid lineage (B-cells, CD4+ T-cells, CD8+ T-cells and lymphoblastoid cell lines) and in two datasets from the myeloid lineage (monocytes and neutrophils). Compared to generic *cis*-eQTLs, large effect sizes were observed for neutrophil-mediated *cis*-eQTLs in myeloid lineage cell types, and small effect sizes in the lymphoid datasets. Conversely, lymphoid-mediated *cis*-eQTL effects had large effect sizes specifically in the lymphoid lineage datasets, while having smaller effect sizes in myeloid lineage datasets. These results indicate that our method is able to reliably predict whether a specific *cis*-eQTL is mediated by cell type. B) Comparison between average gene expression levels between different purified cell type eQTL datasets shows that neutrophil mediated *cis*-eQTLs have, on average a lower expression in cell types derived from the lymphoid lineage, and a high expression in myeloid cell types, while the opposite is true for lymphocyte mediated *cis*-eQTLs.

Myeloid and lymphoid blood cell types provide crucial immunological functions. Therefore, we assessed five immune-related diseases for which genome-wide association studies previously identified at least 20 loci with a *cis*-eQTL in our meta-analysis. We observed a significant enrichment only for Crohn's disease (CD), (binomial test, one-tailed P = 0.002, Supplementary Table 4): out of 49 unique CD-associated SNPs showing a *cis*-eQTL effect, 11 (22%) were neutrophil-mediated. These 11 SNPs affect the expression of 14 unique genes (ordered by size of interaction effect: *IL18RAP, CPEB4, RP11-514O12.4, RNASET2, NOD2, CISD1, LGALS9, AC034220.3, SLC22A4, HOTAIRM2, ZGPAT, LIME1, SLC2A4RG, and PLCL1*). CD is a chronic inflammatory disease of the intestinal tract, and neutrophils are essential for killing microbes that translocate through the mucosal layer of the intestine. The mucosal layer is affected in CD, but also in monogenic diseases with *neutropenia* and defects in phagocyte bacterial killing, such as chronic granulomatous disease, glycogen storage disease type I, and congenital neutropenia, leading to various CD phenotypes (Uhlig 2013). In addition, pharmacological interventions for the treatment of CD have been developed to specifically target neutrophils, including Sagramostim (Korzenik et al. 2005) and Natalizumab (Ghosh et al. 2003). Our new analysis shows clear neutrophil-mediated eQTL effects for many of the known CD genes, including the archetypal *NOD2* gene, and our results provide deeper insight into the role of neutrophils in CD pathogenesis.

Large sample sizes are essential in order to find cell-type-mediated *cis*-eQTLs (Figure 4): when we repeat our study on fewer samples (ascertained by systematically excluding more cohorts from our study), the number of significant cell-type-mediated eQTLs decreased rapidly. This was particularly important for the lymphoid-mediated *cis*-eQTLs, because myeloid cells are approximately twice as abundant as lymphoid cells in whole blood. Consequently, detecting lymphoid-mediated *cis*-eQTLs is more challenging than detecting neutrophil-specific *cis*-eQTLs. As whole blood eQTL data is easily collected, we were able to gather a sufficient sample size in order to detect cell-type-mediated or -specific associations without requiring the actual purification of cell types.

**Figure 4.**
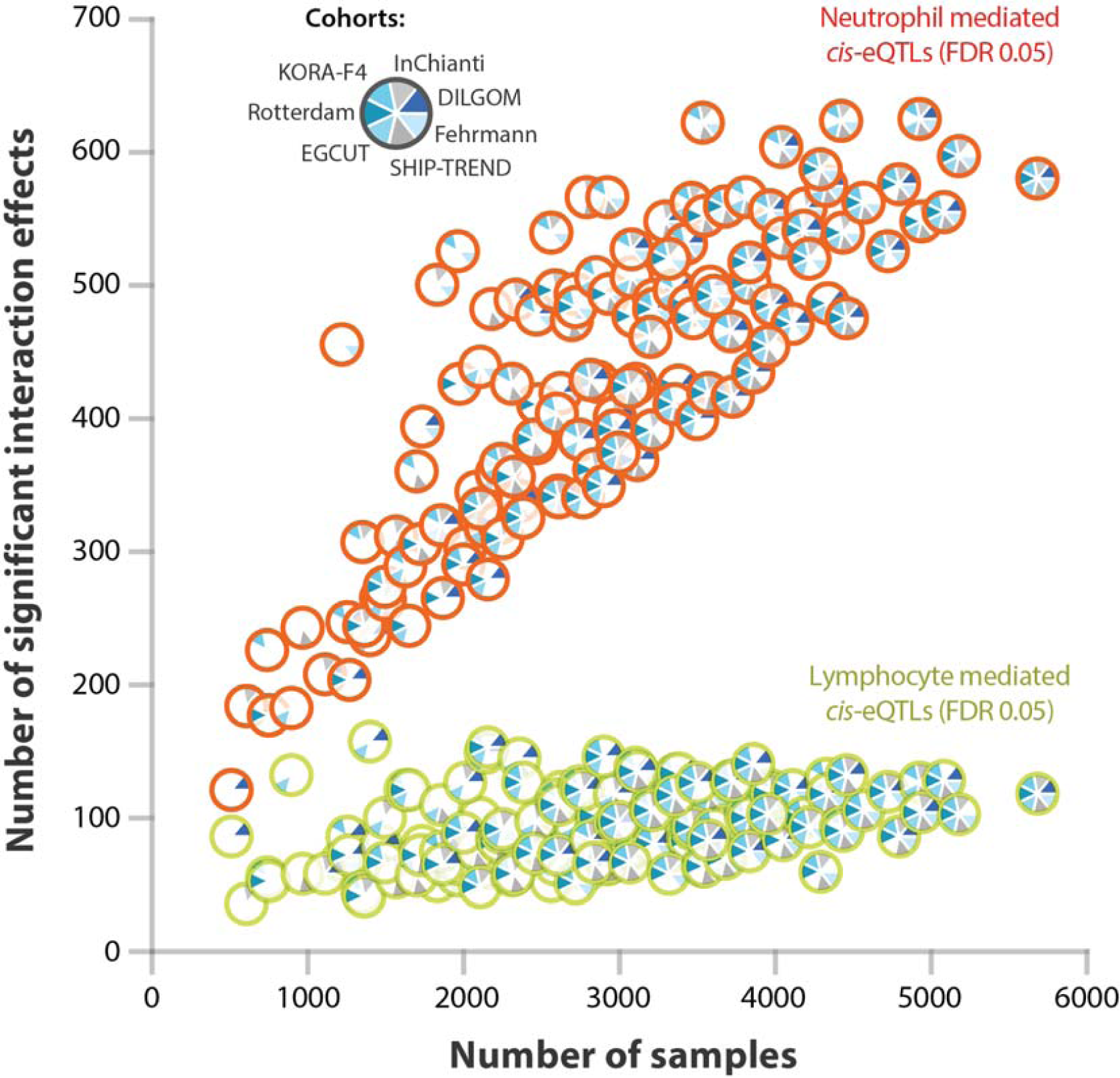
Effect of sample size on power to detect cell type specific *cis*-eQTLs. We systematically excluded datasets from our meta-analysis in order to determine the effect of sample size on our ability to detect significant interaction effects. The number of significant interaction effects was rapidly reduced when the sample size was decreased (the number of unique significant probes given a Bonferroni corrected P-value < 8.1 × 10^−6^ is shown). In general, due to their low abundance in whole blood, lymphoid-mediated *cis*-eQTL effects are harder to detect than neutrophil-mediated *cis*-eQTL effects.

## Discussion

Here we have shown that it is possible to infer in which blood cell-types *cis*-eQTLs are operating from a whole blood dataset. Cell-type proportions were predicted and subsequently used in a G x E interaction model. Hundreds of *cis*-eQTLs showed stronger effects in myeloid than lymphoid cell-types and vice versa.

These results were replicated in 6 individual purified cell-type eQTL datasets (two reflecting the myeloid and four reflecting the lymphoid lineage). This indicates our G x E analysis provides important additional biological insights for many SNPs that have previously been found to be associated with complex (molecular) traits.

Here, we concentrated on identifying *cis*-eQTLs that are preferentially operating in either myeloid or lymphoid cell-types. We did not attempt to assess this for specialized cell-types within the myeloid or lymphoid lineage. However, this is possible if cell-counts are available for these cell-types, or if these cell-counts can be predicted by using a proxy for those cell-counts. As such, identification of cell-type mediated eQTLs for previously unstudied cell-types is possible, without the need to generate new data. However, it should be noted that these individual cell-types typically have a rather low abundance within whole blood (e.g. natural killer cells only comprise ∼2% of all circulating white blood cells). As a consequence, in order to have sufficient statistical power to identify eQTLs that are mediated by these cell-types, very large whole blood eQTL sample-sizes are required, specific cell types should be variable between individuals (which is analogous to the difference in the number of identified lymphoid mediated *cis*-eQTLs, as compared to the number of neutrophil mediated *cis*-eQTLs, which is likely caused by their difference in abundance in whole blood).

We confined our analyses to a subset of *cis*-eQTLs for which we had previously identified a main effect in whole peripheral blood (Westra et al. 2013): for each *cis*-eQTL gene, we only studied the most significantly associated SNP. Considering that for many *cis*-eQTLs multiple, unlinked SNPs exist that independently affect the gene expression levels, it is possible that we have missed myeloid or lymphoid mediation of these secondary *cis*-EQTLs.

We anticipate that the (pending) availability of large RNA-seq based eQTL datasets, statistical power to identify cell-type mediated eQTLs will improve: since RNA-seq enables very accurate gene expression level quantification and is not limited to a set of preselected probes that interrogate well known genes (as is the case for microarrays), the detection of genes that can serve as reliable proxies for individual cell-types will improve. Using RNA-seq data, it is also possible to assess whether SNPs that affect the expression of non-coding transcripts, affect splicing (Lappalainen et al. 2013) or result in alternative polyadenylation (Zhernakova et al. 2013a) are mediated by specific cell-types.

Although we applied our method to whole blood gene expression data, our method can be applied to any tissue, alleviating the need to sort cells or to perform laser capture micro dissection. The only prerequisite for our method is the availability of a relatively small training dataset with cell count measurements in order to develop a reliable proxy for cell count measurements. Since the number of such training datasets is rapidly increasing and meta-analyses have proven successful (Westra et al. 2013; Fehrmann et al. 2011), our approach provides a cost-effective way to identify cell-type-mediated or -specific associations, and it is likely to reveal major biological insights.

## Methods

### Setup of study

This eQTL meta-analysis is based on gene expression intensities measured in whole blood samples. RNA was isolated with either PAXgene Tubes (Becton Dickinson and Co., Franklin Lakes, NJ, USA) or Tempus Tubes (Life Technologies). To measure gene expression levels, Illumina Whole-Genome Expression Beadchips were used (HT12-v3 and HT12-v4 arrays, Illumina Inc., San Diego, USA). Although different identifiers are used across these different platforms, many probe sequences are identical. Meta-analysis could thus be performed if probe-sequences were equal across platforms. Integration of these probe sequences was performed as described before (Westra et al. 2013). Genotypes were harmonized using HapMap2-based imputation using the Central European population (The International HapMap Consortium 2003). In total, the eQTL genotype x environment interaction meta-analysis was performed on seven independent cohorts, comprising a total of 5,863 unrelated individuals. Mix-ups between gene expression samples and genotype samples were corrected using *MixupMapper* (Westra et al. 2011). Gene expression normalization was performed as described before (Westra et al. 2013), removing up to 40 principal components (PCs). Additionally, we corrected for possible confounding factors due to arrays of poor RNA quality, by correlating the sample gene expression measurements against the first PC that was determined from the sample correlation matrix. Samples with a correlation < 0.9 were removed from further analysis.

### Discovery cohorts

#### Fehrmann

The Fehrmann dataset consists of whole peripheral blood samples of 1,240 unrelated individuals from the UK and the Netherlands (Fehrmann et al. 2011; Dubois et al. 2010). Some of these individuals are patients, while others are healthy controls. Individuals were genotyped using Illumina HumanHap300, HumanHap370 or the 610 Quad platform. Genotypes were imputed using Impute v2 (Howie et al. 2009), using the phased genotypes of the CEU subpopulation of HapMap2 release 24 as reference (The International HapMap Consortium 2003). RNA levels were quantified using the HT12v3 platform (N = 1,240), as has been described before (Fehrmann et al. 2011). As sample mix-up correction was performed prior to the participation in this study, the total number of samples with both genotype and gene-expression data, was equal to 1,240. After removing the gene expression samples which showed a poor RNA quality (correlation with PC1 < 0.9), 1,220 samples remained.

#### SHIP-TREND

SHIP (Study of Health in Pomerania, Northeastern Germany) is a population-based project consisting of two independent cohorts, SHIP and SHIP-TREND. The study design of SHIP has been previously described in detail (Völzke et al. 2011). For this eQTL analysis, a part of the the SHIP-TREND cohort was used. The SHIP-TREND probands (N = 986) were genotyped using the Illumina HumanOmni2.5-Quad arrays. Genotypes were imputed to HapMap v2 (The International HapMap Consortium 2003) using IMPUTE (Howie et al. 2009). RNA was prepared from whole blood collected under fasting conditions in PAXgene tubes (Becton Dickinson) using the PAXgene Blood miRNA Kit (Qiagen, Hilden, Germany). For SHIP-TREND this was done on a QIAcube according to protocols provided by the manufacturer (Qiagen). RNA was amplified (Ambion TotalPrep RNA), and hybridized to Illumina HumanHT-12 v3 Expression BeadChip. After sample mix-up correction both imputed genotypes and whole-blood gene expression data were available for a total of 963 SHIP-TREND samples. No samples had to be removed due to poor RNA quality (correlation with PC1 < 0.9).

#### Rotterdam Study

The Rotterdam Study (RS) is a large prospective, population-based cohort study in the district of Rotterdam, the Netherlands, investigating the prevalence, incidence, and risk factors of various chronic disabling diseases among elderly Caucasians aged 45 years and over. The initial cohort, named the Rotterdam Study I (or RS-I) was started in 1989, and consisted of 7,983 persons aged 55 years or over, living in the well-defined Ommoord district. In 1999, a second cohort, named the Rotterdam Study II (or RS-II) was started and consisted of 3,011 participants who had reached the age of 55 years since 1989 or who had moved into the study district. In 2006, the cohort was further extended with 3,932 subjects who were aged 45 years or over; this is called the Rotterdam Study III (RS-III). The Rotterdam Study has been described in detail (Hofman et al. 2011, 2013). Informed consent was obtained from each participant, and the medical ethics committee of the Erasmus Medical Center Rotterdam approved the study.

For this eQTL analysis, we used the RS-III cohort was used. The RS participants (n = 3,054) were genotyped using the Illumina 610 K quad arrays, and genotypes were imputed using MACH (Li et al. 2010) with the HapMap CEU Phase 2 genotypes (release #22, build 36) as a reference (The International HapMap Consortium 2003). Whole blood of 768 samples was collected (PAXgene Tubes, Becton Dickinson) and total RNA was isolated (PAXgene Blood RNA kit, Qiagen). RNA was amplified, labeled (Ambion TotalPrep RNA), and hybridized to the Illumina Whole-Genome Expression Beadchips (Human HT-12v4). The total number of RS-III samples with both imputed genotypes and whole-genome expression data is equal to 768 (before sample mix-up correction). After sample mix-up correction, 762 samples remained. After removing the gene expression samples which showed a poor RNA quality (correlation with PC1 < 0.9), 755 samples remained.

#### EGCUT

The Estonian Gene Expression Cohort (Metspalu 2004) is composed of 899 samples (Mean age 37 [16.6] years; 50% females) from the Estonian Genome Center, University of Tartu (EGCUT) biobank cohort of 53,000 samples. Genotyping was performed using Illumina Human370CNV arrays (Illumina Inc., San Diego, USA), and imputed using Impute v2 (Howie et al. 2009), using the HapMap CEU phase 2 (The International HapMap Consortium 2003) genotypes (release #24, build 36). Whole peripheral blood RNA samples were collected using the Tempus Blood RNA Tubes (Life Technologies), and RNA was extracted using Tempus Spin RNA Isolation Kit (Life Technologies). Quality was measured by NanoDrop 1000 Spectrophotometer (Thermo Fisher Scientific, DE, USA) and Agilent 2100 Bioanalyzer (Agilent Technologies, CA, USA). Whole-Genome gene-expression levels were obtained by Illumina Human HT12v3 arrays (Illumina Inc, San Diego, CA, USA) according to the manufacturer's protocols. After sample mix-up correction, 8 samples were excluded, and 891 samples remained. We did not have to remove any gene expression samples because of poor RNA quality (correlation with PC1 < 0.9), 891 samples remained.

#### DILGOM

The Finnish study samples included a total of 513 unrelated individuals aged 25–74 years from the Helsinki area, recruited during 2007 as part of the Dietary, Lifestyle, and Genetic determinants of Obesity and Metabolic syndrome (DILGOM) study, an extension of the FINRISK 2007 study (Inouye et al. 2010). Study participants were asked to fast overnight (at least 10 hours) prior to giving a blood sample. DNA was extracted from 10 ml EDTA whole blood samples with salt precipitation method using Autopure (Qiagen GmbH, Hilden, Germany). DNA purity and quantity were assessed with PicoGreen (Invitrogen, Carlsbad, CA, USA) and genotyping used 250 ng of DNA which proceeded on the Illumina 610-Quad SNP array (Illumina Inc., San Diego, CA, USA) using standard protocols. SNPs were imputed with MACH version 1.0.10 (Li et al. 2010) using HapMap2 release 22 (The International HapMap Consortium 2003) as a reference panel. To obtain stabilized total RNA, we used the PAXgene Blood RNA System (PreAnalytiX GMbH, Hombrechtikon, Switzerland). It included collection of 2.5 ml peripheral blood into PAXgene Blood RNA Tubes (Becton Dickinson and Co., Franklin Lakes, NJ, USA) and total RNA extraction with PAXgene Blood RNA Kit (Qiagen GmbH, Hilden, Germany). Protocol recommended by the manufacturer was used. The integrity and quantity of the RNA samples were evaluated with the 2100 Bioanalyzer (Agilent Technologies, Santa Clara, CA, USA). Biotinylated cRNA was produced from 200 ng of total RNA with Ambion Illumina TotalPrep RNA Amplification Kit (Applied Biosystems, Foster City, CA, USA), using the protocol specified by the manufacturer. 750 ng of biotinylated cRNA were hybridized onto Illumina HumanHT-12v3 Expression BeadChips (Illumina Inc., San Diego, CA, USA), using standard protocol. After sample mix-up correction, 509 samples were included for further analysis in this cohort. After removing one gene expression sample which showed a poor RNA quality (correlation with PC1 < 0.9), 508 samples remained.

#### InCHIANTI

InCHIANTI (Tanaka et al. 2009) is a population-based, prospective study in the Chianti area (Tuscany) of Italy. The participants were enrolled in 1998-2000, and were interviewed and examined every three years. Ethical approval was granted by the Instituto Nazionale Riposo e Cura Anziani institutional review board in Italy. Participants gave informed consent. Genome-wide genotyping was performed using the Illumina Infinium HumanHap550 genotyping chip. We used MACH 1.0.16 (Li et al. 2010) to impute using the HapMap r22 build-36 reference panel (The International HapMap Consortium 2003). In the InCHIANTI study, peripheral blood specimens were taken using the PAXgene system (PreAnalytiX GMbH, Hombrechtikon, Switzerland), to preserve transcript expression levels. Samples were collected in 2008/9 (wave 4) from 712 participants and mRNA was extracted using the PAXgene Blood mRNA kit (Qiagen, Crawley, UK) according to the manufacturer's instructions. Whole genome expression profiling of the samples was conducted using the Illumina Human HT-12 v3 microarray (Illumina, San Diego, CA, USA) as previously described (Gibbs et al. 2010). Sample mix-up analysis on 620 samples passing QC and having both genotype and gene-expression data, revealed a total of 9 possible sample mix-ups. The total number of InCHIANTI samples with both imputed genotypes and whole-genome expression data included in this analysis was 611. After removing the gene expression samples which showed a poor RNA quality (correlation with PC1 < 0.9), 606 samples remained.

#### KORA F4

KORA F4 (Cooperative Heath Research in the Region of Augsburg, Southern Germany) is a follow-up survey (2006-2008) of the population-based KORA S4 survey that was conducted in the region in 1999-2001. The expression analysis in this study was based on whole blood samples of the KORA F4 participants aged 62 to 81 year (Rathmann et al. 2009; Mehta et al. 2013). RNA was isolated from whole blood using PAXgene Blood miRNA Kit (Qiagen, Hilden, Germany). Purity and integrity of the RNA was analyzed using the Agilent Bioanalyzer with the 6000 Nano LabChip reagent set (Agilent Technologies, Germany). RNA was reverse transcribed with TotalPrep-96 RNA Amp Kit (Ambion, Germany) and hybridized to the Illumina HumanHT-12 v3 Expression BeadChip (Mehta et al. 2013). The samples were genotyped on the Affymetrix 6.0 GeneChip array (Marzi et al. 2010). The SNPs were imputed with MACH (v1.0.15) (Li et al. 2010) and the HapMap CEU version 22 (The International HapMap Consortium 2003) was used as reference population for calling and imputation. Altogether there were 740 samples with gene expression and genotype data available for analysis. We did not have to remove any gene expression samples because of poor RNA quality (correlation with PC1 < 0.9).

### Replication cohorts

#### Stranger LCL

The Stranger Lymphoblastoid Cell Line (LCL) dataset consists of 608 individuals from HapMap3 (Stranger et al. 2012), which were hybridized to Illumina WG6v2 bead chips. As genotypes, we used HapMap3 release 2 and gene expression measurements were normalized per population, using log_2_ transformation, quantile normalization, and principal component (PC) correction. PC correction was limited to 10 PCs because of the small sample size of each individual population. The LCL *cis*-eQTLs were subsequently meta-analyzed over all populations.

#### CD4+ and CD8+ T-cells

Genotypes of this replication cohort are part of the EGCUT study (which is one of the discovery cohorts). As such, they have been generated using the same protocol and quality control measures.

##### Purification of CD4^+^ and CD8^+^ T cells

Peripheral blood was obtained from healthy donors of the Estonian Genome Center of the University of Tartu. Peripheral blood mononuclear cells (PBMC) were extracted using Ficoll-Paque (GE Healthcare) gradient centrifugation. CD4^+^ T cells and CD8^+^ T cells were extracted from the PBMCs by consecutive positive separation using microbeads (CD4+ #130-045-101; CD8+ #130-045-201) and AutoMACS technology (Miltenyi Biotec) according to the manufacturer's protocol. The study was approved by the Ethics Review Committee on Human Research of the University of Tartu, and all of the participants have signed a written informed consent.

##### RNA extraction, labelling and hybridization

RNA was extracted using the miRNeasy Mini Kit combined with a recommended RNase-free DNase I treatment (both from Qiagen) according to the manufacturer's protocol. The RNA was labeled and amplified using the TargetAmp Nano Labeling Kit for Illumina Expression BeadChip (Epicentre Biotechnologies) with SuperScript III Reverse Transcriptase (Life Technologies) according to the manufacturer's protocol, followed by purification with the RNeasy MinElute Cleanup Kit (Qiagen). The labelled RNA samples were hybridized to HumanHT-12 v4 Expression BeadChips (Illumina) according to the manufacturer's instructions. The final number of individuals for this study was 309.

#### Singapore Chinese functional genomics cohort

The Singapore Chinese cohort used for the neutrophil specific eQTL dataset is part of a larger epidemiological cohort described previously (Andiappan et al. 2011). The individuals gave fresh whole blood which was used to isolate neutrophils and frozen down immediately in Trizol to −80° C then extracted from this pure neutrophil population using the Ambion RNA exaction kit. The extracted RNA was then hybridized onto Illumina HumanHT- 12 whole-genome gene expression chips. To avoid batch effects the RNA samples were randomly placed onto Illumina HumanHT-12 arrays such that each chip contains a number of samples from the various RNA extraction batches. Illumina microarray data was then normalized in Genome Studio using quantile normalization with no background subtraction. The log_2_ transformed expression values were then used for subsequent analysis. Additionally whole genome genotyping was done on these individuals using the Human Illumina Omni 5M chip. Samples were then checked for any pair of samples identified as first-degree relatives, and if found, these were removed. SNPs which were monomorphic in the population and those, which failed a call rate of 95% were also removed. This resulted in a total of 4.28 million SNPs with a good call rate that was taken forward for further statistical analysis. Thus, this batch of 114 samples was used to validate that the predicted eQTLs were indeed neutrophil specific.

##### Statistical analysis

Each significant SNP-probe pair in the discovery cohort was analyzed for significance in the validation cohort using a linear regression model. Array address IDs were matched to Illumina probe IDs using the annotation file HumanHT-12 v3. Linear regression was performed using the python function linregress from the library SciPy, coding the genotypes based on allele counts (0, 2 for homozygous genotypes and 1 for heterozygous genotypes). Samples with missing genotypes for a particular SNP were excluded from the analysis of the corresponding SNP-probe pair. 1000 genomes Illumina SNP IDs were mapped to rs IDs based on their annotated position on the chromosome.

#### Oxford

Oxford cell-specific eQTL analysis has been described previously (Fairfax et al. 2012). In the initial analysis peripheral blood mononuclear cell fractions were purified from 50 ml of freshly collected EDTA anti-coagulated blood from 288 healthy European volunteers using Ficoll gradients. CD14+ monocytes and CD19+ B-cells were subsequently positively selected from this fraction using magnetic beads (MACS, Miltenyi-Bitotec, Bergisch Gladbach, Germany) with all steps performed on ice as per the manufacterer's protocol. Individuals were genotyped at 730,525 markers using Illumina OmniExpress Beadchips and, after controlling for population outliers, 283 individuals were used in the final analysis. Genotypes were imputed using the CEU panel of HapMap2 release 24 (The International HapMap Consortium 2003) using BEAGLE v2 (Browning and Browning 2009). The final sample size for B-cells was 282 and the final sample size for monocytes was 283.

#### Gene expression normalization

Each cohort performed gene expression normalization individually: gene expression data was quantile normalized to the median distribution then log_2_ transformed. The probe and sample means were centered to zero. Gene expression data was then corrected for possible population structure by removing four multi-dimensional scaling components (MDS components obtained from the genotype data using PLINK) using linear regression. Because normalized gene-expression data still contains large amounts of non-genetic variation (Schurmann et al. 2012; Fehrmann et al. 2011), principal component analysis (PCA) was performed on the sample correlation matrix, and up to 40 principal components (PCs) were then removed from the gene expression data using linear regression (Westra et al. 2013).

In order to improve statistical power to detect cell-type mediated eQTLs, we corrected the gene expression for technical and batch effects (here we applied principal component analysis and removed per cohort the 40 strongest principal components that affect gene expression). Such procedures are commonly used when conducting *cis*-eQTL mapping (Dubois et al. 2010; Fehrmann et al. 2011; Fu et al. 2012; Zhernakova et al. 2013b; Westra et al. 2013; Lappalainen et al. 2013). To minimize the amount of genetic variation removed by this procedure, we performed QTL mapping for each principal component, to ascertain whether genetic variants could be detected that affected the PC. If such an effect was detected, we did not correct the gene expression data for that particular PC (Westra et al. 2013). We chose to remove 40 PCs based on our previous study results, which suggested that this was the optimum for detecting eQTLs (Westra et al. 2013). The PC-corrected gene expression data was then used as the predicted variable in our model.

### Creating a proxy for neutrophil cell percentage from gene expression data

To be able to determine whether a *cis*-eQTL is mediated by neutrophils, we reasoned that such a *cis*-eQTL would show a larger effect size in individuals with a higher percentage of neutrophils than in individuals with a lower percentage. However, this required the percentage of neutrophils in whole blood to be known, and cell-type percentage measurements were not available for all of the cohorts. We therefore created a proxy phenotype that reflected the actual neutrophil percentage that would also be applicable to datasets without neutrophil percentage measurements. In the EGCUT dataset, we first quantile normalized and log_2_ transformed the raw expression data. We then correlated the gene expression levels of individual probes with the neutrophil percentage, and selected 58 gene expression probes showing a high positive correlation (r^2^ > 0.3).

In each independent cohort, we corrected for possible confounding factors due to arrays with poor RNA quality, by correlating the sample gene expression measurements against the first PC determined from the sample correlation matrix. Only samples with a high correlation (r ≥ 0.9) were included in further analyses. Then, for each cohort, we calculated a correlation matrix for the neutrophil proxy probes (the probes selected from the EGCUT cohort). The gene expression data used was quantile normalized, log_2_ transformed and corrected for MDS components. Applying PCA to the correlation matrix, we then obtained PCs that described the variation among the probes selected from the EGCUT cohort. As the first PC (PC1) contributes the largest amount of variation, we considered PC1 as a proxy-phenotype for the cell type percentages.

### Determining cell-type mediation using an interaction model

Considering the overlap between the cohorts in this study and our previous study, we limited our analysis to the 13,124 *cis*-eQTLs having a significant effect (false discovery rate, FDR < 0.05) in our previous study (Westra et al. 2013). This included 8,228 unique Illumina HT12v3 probes and 10,260 unique SNPs (7,674 SNPs that showed the strongest effect per probe, and 2,586 SNPs previously associated with complex traits and diseases, as reported in the Catalog of Published Genome-Wide Association Studies (Hindorff et al. 2009), on 23^rd^ September, 2013).

We defined the model for single marker *cis*-eQTL mapping as follows:

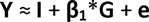

where **Y** is the gene expression of the gene, **β**_1_ is the slope of the linear model, **G** is the genotype, **I** is the intercept with the y-axis, and **e** is the general error term for any residual variation not explained by the rest of the model.

We then extended the typical linear model for single marker *cis*-eQTL mapping to include a covariate as an independent variable, and captured the interaction between the genotype and the covariate using an interaction term:

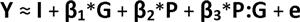

where P (cell-type proxy) is the covariate, and P:G is the interaction term between the covariate and the genotype. We used gene expression data corrected for 40 PCs as the predicted variable (**Y**). The interaction terms were then meta-analyzed over all cohorts using a Z-score method, weighted for the sample size (Whitlock 2005).

### Multiple testing correction

Since the gene-expression data has a correlated structure (i.e. co-expressed genes) and the genotype data also has a correlated structure (i.e. linkage disequilibrium between SNPs), a Bonferroni correction would be overly stringent. We therefore first estimated the effective number of uncorrelated tests by using permuted eQTL results from our previous *cis*-eQTL meta-analysis (Westra et al. 2013). The most significant P-value in these permutations was 8.15 × 10^−5^, when averaged over all permutations. As such, the number of effective tests = 0.5/8.15 × 10^−5^ ≈ 6134, which is approximately half the number of correlated *cis*-eQTL tests that we conducted (=13,124). Next, we controlled the FDR at 0.05 for the interaction analysis: for a given P-value threshold in our interaction analysis, we calculated the number of expected results (given the number of effective tests and a uniform distribution) and determined the observed number of eQTLs that were below the given P-value threshold (FDR = number of expected p-values below threshold/number of observed p-values below threshold). At an FDR of 0.05, our nominal p-value threshold was 0.009 (corresponding to an absolute interaction effect Z-score of 2.61).

### Cell-type specific *cis*-eQTLs and disease

For each trait in the GWAS catalog, we pruned all SNPs with a GWAS association P-value below 5 × 10^−8^, using an r^2^ threshold of 0.2. We only considered traits that had more than 20 significant eQTL SNPs after pruning (irrespective of cell-type mediation). Then, we determined the proportion of pruned neutrophil-mediated *cis*-eQTLs for the trait relative to all the neutrophil-mediated *cis*-eQTLs. The difference between both proportions was then tested using a binomial test.

## Data Access

The source code and documentation for this type of analysis are available as part of the eQTL meta-analysis pipeline at https://github.com/molgenis/systemsgenetics

Summary results are available from http://www.genenetwork.nl/celltype

## Accession numbers

Discovery cohorts: Fehrmann (GSE 20142), SHIP-TREND (GSE 36382), Rotterdam Study (GSE 33828), EGCUT (GSE 48348), DILGOM (E-TABM-1036), InCHIANTI (GSE 48152), KORA F4 (E-MTAB-1708).

Replication Cohorts: Stranger (E-MTAB-264), Oxford (E-MTAB-945).

## DILGOM

J.K. and S.R. were supported by funds from The European Community's Seventh Framework Programme (FP7/2007-2013) BioSHaRE, grant agreement 261433, S.R. was supported by funds from The European Community's Seventh Framework Programme (FP7/2007-2013) ENGAGE Consortium, grant agreement HEALTH-F4-2007-201413”, the Academy of Finland Center of Excellence in Complex Disease Genetics (grants 213506 and 129680), Academy of Finland (grant 251217), the Finnish foundation for Cardiovascular Research and the Sigrid Juselius Foundation. V.S. was supported by the Academy of Finland, grant number 139635 and Finnish Foundation for Cardiovascular Research. MP was partly financially supported for this work by the Finnish Academy SALVE program ‘‘Pubgensense’’ 129322 and by grants from the Finnish Foundation for Cardiovascular Research. The DILGOM-study was supported by the Academy of Finland, grant # 118065.

## SHIP-TREND

SHIP is part of the Community Medicine Research net of the University of Greifswald, Germany, which is funded by the Federal Ministry of Education and Research (grants no. 01ZZ9603, 01ZZ0103, and 01ZZ0403), the Deutsche Forschungsgemeinschaft (DFG GRK840-D2), the Ministry of Cultural Affairs as well as the Social Ministry of the Federal State of Mecklenburg-West Pomerania, and the network ‘Greifswald Approach to Individualized Medicine (GANI_MED)’ funded by the Federal Ministry of Education and Research (grant 03IS2061A). Genome-wide data have been supported by the Federal Ministry of Education and Research (grant no. 03ZIK012) and a joint grant from Siemens Healthcare, Erlangen, Germany and the Federal State of Mecklenburg, West Pomerania. Whole-body MR imaging was supported by a joint grant from Siemens Healthcare, Erlangen, Germany and the Federal State of Mecklenburg West Pomerania. The University of Greifswald is a member of the ‘Center of Knowledge Interchange’ program of the Siemens AG and the Caché Campus program of the InterSystems GmbH. The SHIP authors thank Mario Stanke for the opportunity to use his Server Cluster for the SNP imputation.

## EGCUT

EGCUT received financing by FP7 grants (201413, 245536), also received targeted financing from the Estonian Government (SF0180142s08) and direct funding from the Ministries of Research and Science and Social Affairs. EGCUT studies are funded by the University of Tartu in the framework of the Center of Translational Genomics and by the European Union through the European Regional Development Fund, in the framework of the Centre of Excellence in Genomics. We thank EGCUT personnel, especially Ms. M. Hass and Mr V. Soo. EGCUT data analyses were carried out in part in the High Performance Computing Center of the University of Tartu.

## Rotterdam Study

We thank Pascal Arp, Mila Jhamai, Marijn Verkerk, Lizbeth Herrera, and Marjolein Peters for their help in creating the GWAS database; Karol Estrada and Maksim Struchalin for their support in creation and analysis of imputed data; Tobias A. Knoch, Anis Abuseiris, Karol Estrada, and Rob de Graaf as well as their institutions, the Erasmus GRID Office, Erasmus MC Rotterdam, The Netherlands, and especially the national German MediGRID and Services@MediGRID part of the German D-Grid, both funded by the German Bundesministerium fuer Forschung und Technology under grants #01 AK 803 A-H and # 01 IG 07015 G for access to their grid resources. The authors thank the study participants and staff from the Rotterdam Study, the participating general practitioners and the pharmacists.

The Rotterdam Study was funded by the European Commission (HEALTH-F2-2008-201865, GEFOS; HEALTH-F2-2008 35627, TREAT-OA 200800), the Netherlands Organization of Scientific Research NWO Investments (nos 175.010.2005.011, 911-03-012), the Research Institute for Diseases in the Elderly (014-93-015; RIDE2), the Netherlands Genomics Initiative (NGI)/Netherlands Consortium for Healthy Aging (NCHA) (project nr. 050-060-810), an NWO VIDI grant (#917103521).

The Rotterdam Study is funded by Erasmus Medical Center and Erasmus University, Rotterdam, Netherlands Organization for Health Research and Development (ZonMw), the Research Institute for Diseases in the Elderly (RIDE), the Ministry of Education, Culture and Science, the Ministry for Health, Welfare and Sports, the European Commission (DG XII), and the Municipality of Rotterdam.

## Fehrmann

L.F.,H-J.W.: This study was supported by grants from the Celiac Disease Consortium (an innovative cluster approved by the Netherlands Genomics Initiative and partly funded by the Dutch Government (grant BSIK03009), the Netherlands Organization for Scientific Research (NWO-VICI grant 918.66.620, NWO-VENI grant 916.10.135 to L.F.), the Dutch Digestive Disease Foundation (MLDS WO11-30), and a Horizon Breakthrough grant from the Netherlands Genomics Initiative (grant 92519031 to L.F.). This project was supported by the Prinses Beatrix Fonds, VSB fonds, H. Kersten and M. Kersten (Kersten Foundation), The Netherlands ALS Foundation, and J.R. van Dijk and the Adessium Foundation. The research leading to these results has received funding from the European Community's Health Seventh Framework Programme (FP7/2007-2013) under grant agreement 259867. We especially thank Jackie Senior and Kate McIntyre for critically reading the manuscript. This study was supported by the BBMRI NL Functional Genomics Project. Funding for the project was provided by the Netherlands Organization for Scientific Research under award number 184021007, dated July 9, 2009 and made available as a Rainbow Project of the Biobanking and Biomolecular Research Infrastructure Netherlands (BBMRI–NL). D.A. was supported by the Centre for BioSystems Genomics (CBSG) and the Netherlands Consortium of Systems Biology (NCSB), both of which are part of the Netherlands Genomics Initiative/Netherlands Organisation for Scientific Research.

## InCHIANTI

InCHIANTI was supported by the Wellcome Trust 083270/Z/07/Z. The InCHIANTI study was supported by contract funding from the U.S. National Institute on Aging (NIA), and the research was supported in part by the Intramural Research Program, NIA, and National Institute of Health (NIH). A.R.W. was supported by the Peninsula NIHR Clinical Research Facility. Funding to pay the Open Access publication charges for this article was provided by the Wellcome Trust.

## KORA F4

The KORA authors acknowledge the contributions of Peter Lichtner, Gertrud Eckstein, Guido Fischer, Norman Klopp, Nicole Spada, and all members of the Helmholtz Zentrum München genotyping staff for generating the SNP data and Katja Junghans and Anne Löschner (Helmholtz Zentrum München) for generating gene expression data from both KORA and SHIP-TREND samples.

The KORA research platform and the KORA Augsburg studies are financed by the Helmholtz Zentrum München, German Research Center for Environmental Health, which is funded by the BMBF and by the State of Bavaria. We thank the field staff in Augsburg who were involved in the studies. The German Diabetes Center is funded by the German Federal Ministry of Health and the Ministry of School, Science and Research of the State of North-Rhine-Westphalia. The Diabetes Cohort Study was funded by a German Research Foundation project grant to W.R. (DFG; RA 459/2-1). This study was supported in part by a grant from the BMBF to the German Center for Diabetes Research (DZD e.V.), by the DZHK (Deutsches Zentrum für Herz-Kreislauf-Forschung – German Centre for Cardiovascular Research) and by the BMBF funded Systems Biology of Metabotypes grant (SysMBo#0315494A). Additional support was given by the BMBF (National Genome Research Network NGFNplus Atherogenomics, 01GS0834) and the Leibniz Association (WGL Pakt für Forschung und Innovation). We thank Maren Carstensen, Gabi Gornitzka and Astrid Hoffmann (German Diabetes Center) for excellent technical assistance.

## Oxford

This work was supported by the Wellcome Trust (Grants 074318 [J.C.K.], 088891 [B.P.F.], and 075491/Z/04 [core facilities Wellcome Trust Centre for Human Genetics]), the European Research Council under the European Union's Seventh Framework Programme (FP7/2007-2013)/ERC Grant agreement no. 281824 (J.C.K.) and the NIHR Oxford Biomedical Research Centre.

## Singapore Chinese functional genomics cohort

These studies were supported by A*STAR/SIgN core funding, and grants SIgN-06-006, SIgN-08-020 and SIgN-10-029.

## Author contributions

Development of the cell type specific eQTL mapping method: H-J.W., D.A., R.C.J. and L.F.

Computational analysis and interpretation of the results: H-J.W., D.A., T.E., M.J.P., C.S., K. Schramm, J. Kettunen, J. Karjalainen, H.Y., B.P.F., S.K., R.M., B.T., M. Poidinger and R.C.J.

Reviewing and editing of the manuscript: H-J.W., D.A., T.E., M.J.P., C.S., K. Schramm, J.K., A.K.A., Y.L., J.F, M.C., R.K., C.W., S.K., L.M., L.T., P.P., E.R., A.H., A.G.U., F.R., G.H., H.P., T.M., C.H., M.R., H.G., S.R., A.R.W., D.M., L.F., A.B.S., D.G.H, R.M., B.T., M. Poidinger, F.Z., A.L., D.Y.W., O.R., K.S., U.V., J.B.J.M., A.M., R.C.J. and L.F.

Data collection: T.E., H.Y., B.P.F., A.K.A., M. Platteel, L.M., L.T., P.P., E.R., A.H., A.G.U., A.P., R.L., H.P., T.M., C.H., M.R., H.G., M. Perola, S.M., J.C.K., D.Y.W., L.H.v.d.B, J.H.V., O.R., T.M.F., V.S., K. Strauch and A.M.

---

**Supplementary Figure 1 Comparison of effect sizes and effect direction between datasets**

Comparison of interaction effect Z-scores shows a high consistent direction of effect between datasets and with the meta-analysis for those interaction effects significant at FDR < 0.05.

**Supplementary Table 1**

Summary statistics for the interaction analysis.

**Supplementary Table 2**

Results of the interaction analysis.

**Supplementary Table 3**

Summary statistics showing the effect size (correlation coefficient) in each of the tested replication datasets.

**Supplementary Table 4**

Results of the neutrophil mediated *cis*-eQTL disease enrichment analysis.

